# LPMO-oxidized cellulose oligosaccharides evoke immunity in Arabidopsis conferring resistance towards necrotrophic fungus *B. cinerea*

**DOI:** 10.1101/2021.04.28.441599

**Authors:** Marco Zarattini, Massimiliano Corso, Marco Antonio Kadowaki, Antonielle Monclaro, Silvia Magri, Irma Milanese, Sylvie Jolivet, Mariana Ortiz de Godoy, Christian Hermans, Mathilde Fagard, David Cannella

## Abstract

Lytic Polysaccharide Monooxygenases (LPMOs) are powerful redox enzymes able to oxidatively cleave cellulose polymers. Widely conserved across biological kingdoms, LPMOs of the AA9 family are deployed by phytopathogens during necrotrophic attack of plant cell wall. In response, plants have evolved sophisticated mechanisms to sense cell wall damage and thus self-triggering Damage Triggered Immunity (DTI) responses. Here, we show that Arabidopsis plants exposed to LPMO products responds by activating the innate immunity ultimately leading to increased resistance to pathogenic fungus Botrytis cinerea. We demonstrated with microarray hybridization that plants undergo a deep transcriptional reprogramming upon elicitation with AA9 derived cellulose- or cello-oligosaccharides (AA9_COS). To decipher the specific effects of native and oxidized LPMO-generated cello-oligosaccharides, a pairwise comparison with cellobiose, the smallest non-oxidized unit constituting cellulose, is presented. Moreover, we identified two leucine-rich repeat receptor-like kinases, namely STRESS INDUCED FACTOR 2 and 4, playing a crucial role in signaling the AA9_COS-dependent responses such as camalexin production. We observed an increased production of ethylene, jasmonic and salicylic acid hormones, and finally deposition of callose in cell wall. Collectively, our data reveal that LPMOs might play a crucial role in plant-pathogen interactions.

## Introduction

The plant cell wall is a dynamic structure mostly made up of high molecular weight polysaccharides, i.e., cellulose, hemicellulose, pectin, and the heteropolymer lignin^1^. This complex aggregate confers structural integrity and physical protection to plant cells^1^. To overcome the cell wall barrier, phytopathogens have developed an ingenious arsenal of enzymes collectively referred to as cell wall degrading enzymes (CWDEs) ^2^. Thus, the structural cell wall integrity (CWI) is constantly monitored through dedicated molecular sensors and unique signaling mechanisms, including diffusible cell wall-derived molecules, i.e oligosaccharides^3,4,5^.

A novel family of CWDEs, the Lytic Polysaccharide Monooxygenases (LPMOs), was discovered in 2010^6^ and, despite its potential role in plant pathogenicity^7^, its biological significance in plant-pathogen interactions is still overlooked. LPMOs are metalloenzymes, bearing a mono-copper atom in a unique T-shape histidine-brace pocket that catalyzes the oxidative cleavage of 1,4 glycosidic bonds of polysaccharides including cellulose, chitin, starch and xyloglucans^8^. The oxidative cleavage can take place at either C1- or C4-position of the pyranose ring. This cleavage yields a full array of oxidized and native cellulose- or cello-oligosaccharides (COS), i.e. glucose polymers of variable degrees of polymerization (DP from 2 to 10), as well as their C1- or C4-oxidized counterparts^6^. The catalytic mechanism, still under debate, depends on the presence of O_2_ or H_2_O_2_ as electron sink for the redox activity, however an external electron donor is always needed to cleave polysaccharides^9^. During in vitro tests, reductants such as ascorbate or gallate are often used to transfer electrons to LPMO. In addition, lignin-derived phenols^10;11;12^, photoactivated pigments such as chlorophyllin^9,13^, and protein partners like cellobiose dehydrogenases (CDH) ^14^, are all possible electron donors for LPMOs in *vivo* activity^15^

Widely distributed across the entire tree-of-life^16^, the LPMOs are particularly abundant in bacterial and fungal kingdoms^17^. According to the carbohydrate-active enzymes database (www.cazy.org), LPMOs are classified into seven “auxiliary activity” (AA) families: AA9-11 till AA13-16. The genome of several organisms (especially phytopathogenic fungi) features multiple LPMO gene copies. For example, *Fusarium graminearum and Botrytis cinerea* count respectively thirteen and ten AA9 isoforms^18^, a family generally active against cellulose and xyloglucans^8^. So far none LPMO active on plant cell wall polysaccharides were found in plant’s genome.

Plant cells possess at least three partially overlapping layers of defenses defined as Pattern Triggered Immunity (PTI), Damage Triggered Immunity (DTI) and Effector Triggered Immunity (ETI), which constitute the so-called plant immune system^19^. The ETI confers a robust resistance response, as it is initiated following recognition of pathogen virulence effectors (Avr-proteins) by plant cytoplasmic resistance genes (R-genes) ^20^. On the contrary, PTI triggers less powerful resistance responses as compared to ETI, but it provides broad-spectrum protection^19^. PTI is initiated upon recognition of evolutionary-conserved Pathogen-Associated Molecular Patterns (PAMPs), which are sensed by plants through a plethora of plasma membrane-anchored pattern-recognition receptors (PRRs) ^21^. Similarly, plants can self-trigger DTI by recognizing Damage-Associated Molecular Patterns (DAMPs), which comprise cell wall-derived molecules as well as de *novo* synthesized stress-associated peptides^19,22,23^.

Common signaling events underlying PTI/DTI involve Ca^2+^ influx into the cytoplasm^24^, activation of mitogen-activated protein kinase (MAPK) cascades^25^, reactive oxygen species (ROS) accumulation^26^, and an extensive transcriptional reprogramming including expression of transcription factors^27^. During PTI/DTI, plants accumulate signaling hormones such as salicylic acid (SA), jasmonic acid (JA) and ethylene (ET) while several hours after PTI/DTI activation, callose deposition occurs at the cell wall^25,28^. Besides, PTI/DTI triggers the synthesis of a wide range of specialized metabolites including camalexin, an indolic phytoalexin that plays an essential role in plant resistance^29^. Noteworthy, plant defenses are governed by spatial and temporal dynamics^27^. For example, signaling events triggered by PAMPs or DAMPs perception occur within seconds to minutes, whereas transcriptome, cellular and physiological responses are triggered within hours to days.

Thorough studies have investigated the elicitor properties of DAMPs derived from the hydrolysis of polysaccharides such as pectin and hemicellulose, i.e., oligogalacturonides (OGs) ^30,31^ and xyloglucan^32^. More recently, the community started to investigate deeper structures of the plant cell wall. Cellulose-derived molecules, mainly cellobiose^33^ and cellotriose^34^ generated by glycoside hydrolase (GH) enzymes, were demonstrated to behave as DAMPs triggering plant immunity^35^. On the other hand, little is known about the biological implications of oxidative deconstruction of cellulose conducted by AAs enzymes, especially those belonging to the redox family of LPMOs. Although a hypothetical role in plant pathogenesis was speculated^18^, the biological effects of cellulose deconstruction via oxidative mechanism, their perception and the triggered signaling remained so far an elusive idea.

In this study, the TtAA9E from the fungus *Thermothielavioides terrestris* was chosen as a representative member of the AA9 LPMO family, to produce a pool of cello-oligosaccharides comprised of native and their C1- and C4-oxidized counterparts (hereafter named AA9_COS). We highlighted that AA9_COS treatment leads to a strong transcriptomic reprogramming in Arabidopsis, mainly associated to plant immunity. Together with other biotic stress responses, the antimicrobial phytoalexin camalexin markedly accumulated after AA9_COS treatment. Moreover, we identified two LRR-RLK proteins, namely *Stress Induced Factor 2 and 4*, crucial to signal the AA9_COS-dependent responses. Most importantly, a reduction of B. *cinerea* propagation was observed in AA9_COS-treated plants, paving a route for a biotechnological application for these molecules.

## Results

### AA9 genes of *Botrytis cinerea* are expressed during plant infection and encode enzymes with putative C1/C4 regioselectivity as *Tt*AA9E

In this work we obtain oxidized COS from the conversion of cellulose using the TtAA9E enzyme produced by the fungus T. *terrestris* (Fig. 1). This family of enzymes is present on the necrotrophic pathogen B. cinerea, and they are selectively expressed during the infective process in Arabidopsis and none during growth on dextrose media (Supplementary Fig. 1). The TtAA9E enzyme feature a C1/C4 regioselectivity^13,36^ during the oxidative cleavage of the β-1,4-glycosidc bonds when incubated on the highly accessible phosphoric acid swollen cellulose (PASC), that was purposely chosen as the ideal substrate to ensure stable and sufficient amount of AA9_COS. The COS so obtained were purified with molecular filter (3KDa cut-off), the absence of residual peptides was checked with SDS-PAGE (Supplementary Fig 2) and further characterized with High-Performance Anion-Exchange Chromatography with Pulsed Amperometric Detection (HPAEC-PAD) (Fig. 1c). The mixture of cellulose-derived COS presented both native cello-oligosaccharides and their aldonic C1-oxidized forms with various degrees of polymerization (DP2-7) with a slight enrichment towards cellobionic, cellotrionic and cellotetraonic acid compared to their native counterparts. Whereas the C4-oxidized (gemdiol) COS although detected, represented a very minor fraction of the entire mixture (Fig. 1c).

**Fig. 1:**
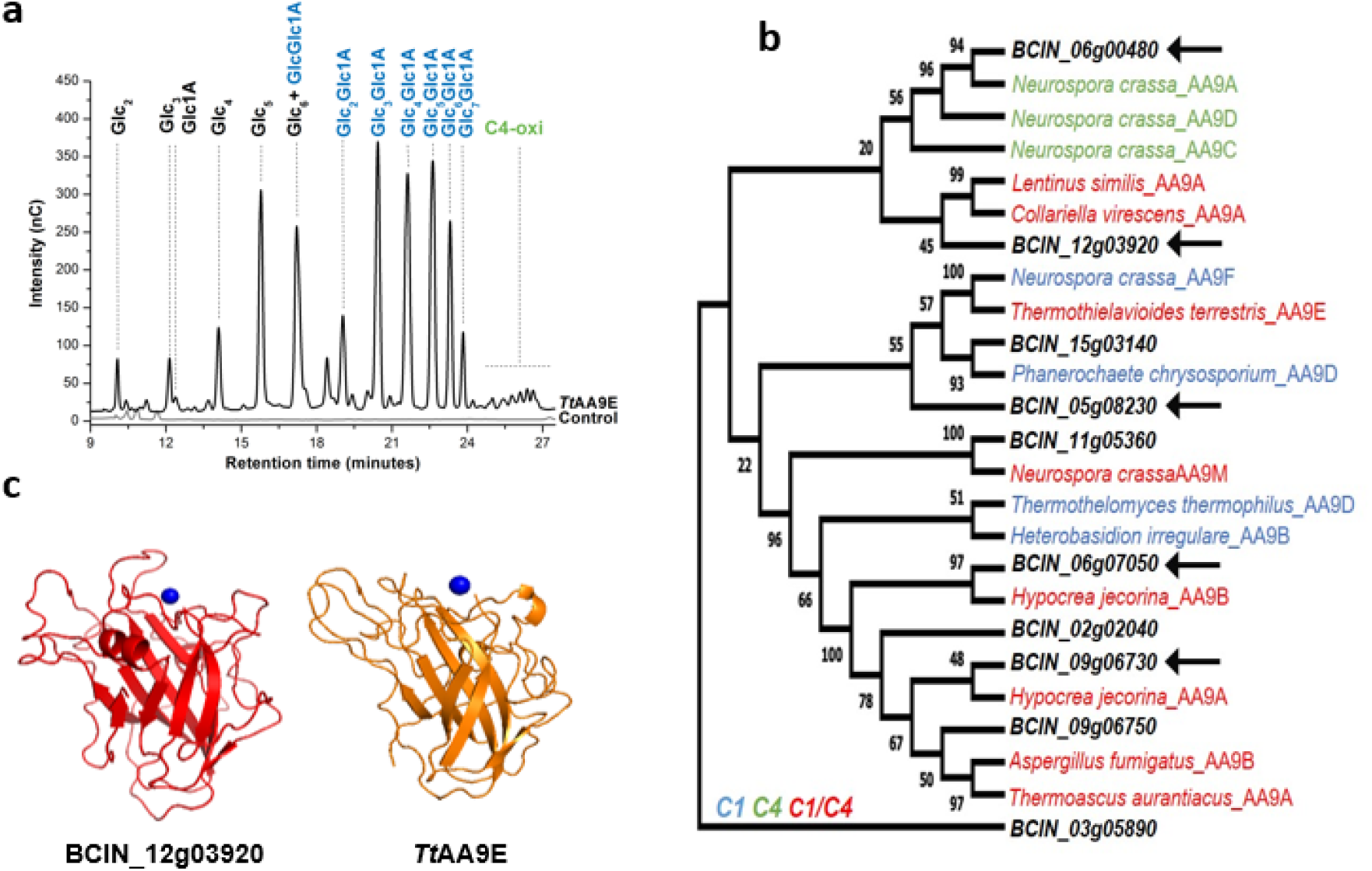
*Bc*AA9s sequence alignment, structural prediction and *Tt*AA9E-derived cello-oligosaccharides characterization. **a** HPAEC-PAD analysis of the products generated by TtAA9E using PASC 0.5% (w/v) as cellulosic substrate. The shown profile is the average of five independent experiments and relative chromatograms. Glc2: cellobiose, Glc3: cellotriose, Glc4: cellotetraose, Glc5: cellopentaose, GlcGlc1A: cellobionic acid, Glc2Glc1A: cellotrionic acid, Glc3Glc1A: cellotetraonic acid, Glc4Glc1A: cellopentaoinic acid, Glc5Glc1A: cellohexaoinic acid, Glc6Glc1A: celloeptaonic acid, Glc7Glc1A: cellooctaonic acid. **b** Sequence-based comparison of *Bc*AA9s with AA9 enzymes biochemically and structurally characterized. The phylogenetic tree was created based on the alignment of catalytic domain. Colours indicated the oxidative regiospecificity: C1, blue; C4, green; C1/C4, red; not characterized, black. The most expressed BcAA9s during *B. cinerea*-Arabidopsis interaction are labelled with arrows. **c** Structural comparison between *Tt*AA9E and BCIN_12g03920 was built using the Swiss-Model Server and the *Tt*AA9E crystal structure obtained from the protein data bank (PDB:3EII). The structural models are shown in ribbon representation and copper ion is shown as a blue sphere.

Interestingly, a similar regioselectivity and structure was predicted from sequence alignment for some *B. cinerea* AA9s (*Bc*AA9) and structurally resolved AA9s. In particular, the BCIN_12g03920 folding prediction shown a highly similarity in structure compared to *Tt*AA9E (Fig. 1b, 1c). BcAA9s were already shown being expressed during necrotrophic growth in crops and fruits^7^, therefore we investigated if also during Arabidopsis infection these could be expressed. The mRNA accumulation of the entire *BcAA9* gene family was measured 48 h after inoculation of Arabidopsis leaves (Supplementary Fig. 1). Five out of ten BcAA9 genes showed substantial overexpression (Fold change > 2) when compared to growth on dextrose medium. Among them BCIN_12g03920 showed particularly high expression *in planta* (Supplementary Fig. 1). The phylogenetic tree in Fig. 1c comparing the *Bc*AA9s catalytic domain sequences with structurally-resolved AA9s enzymes, showed three (BCIN_06g07050, BCIN_09g06730 and BCIN_12g03920) out of the five *in planta*-expressed *Bc*AA9 genes clustering with AA9s enzymes from other species featuring C1/C4-regioselectivity.

### Treatment with LPMO-generated COS protects plants against *B. cinerea*

Upon PTI or DTI activation the plant defence responses follow distinct temporal activation dynamics^27^; therefore, the most appropriate timepoint for the detection of each specific response was chosen accordingly. To evaluate if treatments with AA9_COS conferred resistance toward phytopathogens, as previously demonstrated for pectin-derived OGs^37^, we first performed a dose-response analysis of two defense marker genes 1 h after treatment. The *FLG22-INDUCED RECEPTOR-LIKE KINASE 1 (FRK1)* and *WRKY DNA-BINDING PROTEIN 18* (WRKY18) genes^38^ showed a significant induction when plants were treated with 100 µM AA9_COS (Supplementary Fig. 3). Therefore, rosettes of Arabidopsis plants were drop-treated with 100 µM AA9_COS and inoculated with *B. cinerea* spore suspension (5·10^5^ spores mL^-1^) 24 h after AA9_COS treatment. Besides mock treatment, a solution containing an equimolar concentration of cellobiose was used as a positive control treatment^33^. The *B. cinerea* necrotic lesions sizes were significantly (p < 0.05) reduced by 30% upon AA9_COS treatment when compared with mock-treated leaves (Fig. 2b). These data are in accordance with the *B. cinerea in planta* growth performed three days after infection which highlighted a reduction in fungus growth of approximately 60% (Fig. 2c). On the other hand, a lower protective effect, in terms of lesion size and fungal *in planta* growth, was observed in Arabidopsis plants treated with 100 µM cellobiose (Fig. 2a-c). Interestingly, similar protective effects of AA9_COS treatment against *B. cinerea* were observed in two-month-old tomato plants (*S. lycopersicum* L.; Supplementary Fig. 4a, b).

**Fig 2:**
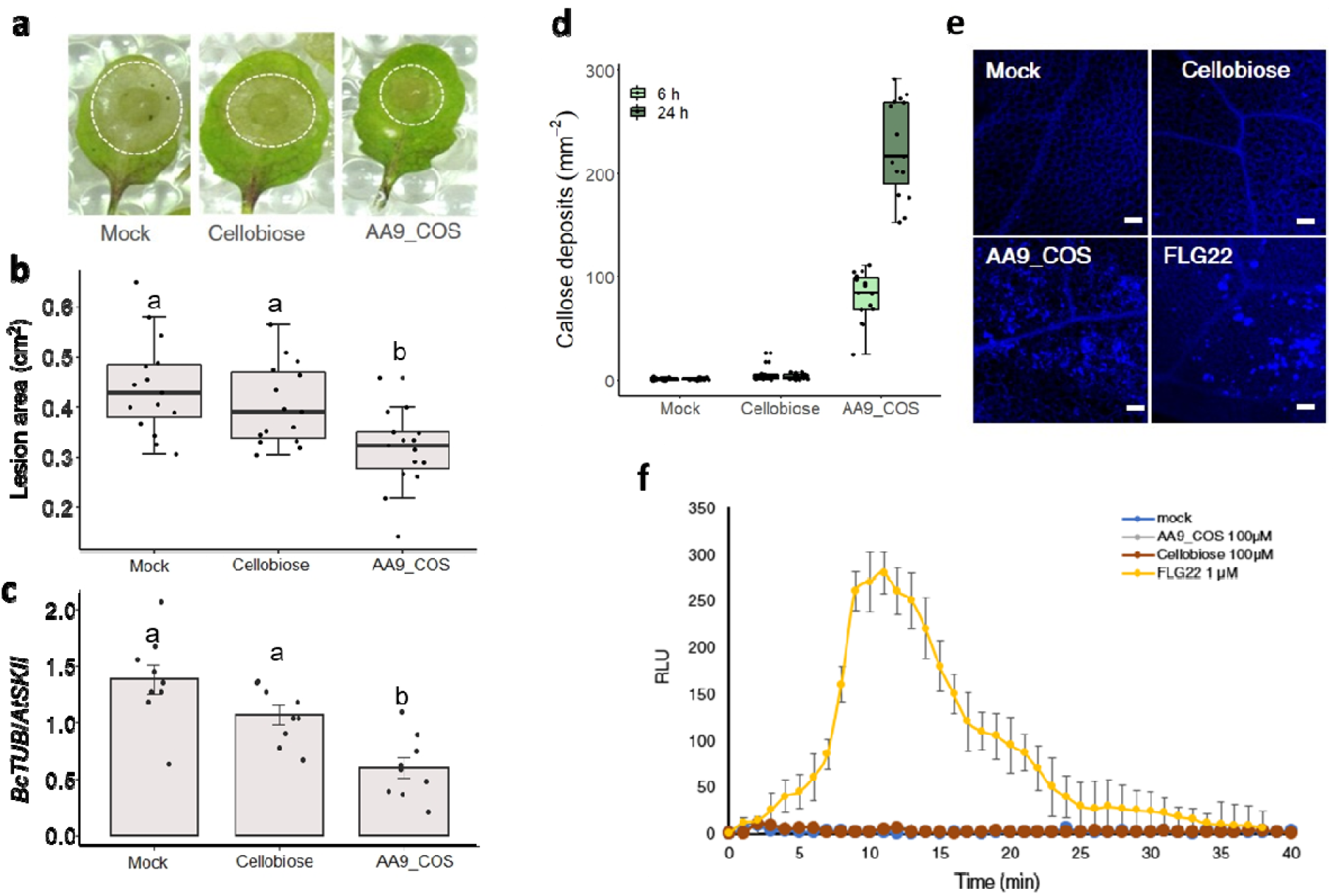
AA9_COS induces DTI responses and protects against *B. cinerea*. **a** Five-week-old Arabidopsis plants were treated with 100 µM cellobiose, 100 µM AA9_COS or mock and 24 h later, detached leaves were spotted with 10 µL spore suspension (5 10^5^ spores mL^-1^). Images were taken three days after infection and representative pictures are reported. **b** Box plots showing median value of the area of necrotic symptoms reported in a. Five leaves from three plants (n = 3) were analyzed. This experiment was repeated three times. **c** The *in-planta* growth of B. cinerea was determined three days after infection by qRT-PCR using housekeeping genes specific for Arabidopsis (*At*SKII) and *B. cinerea* (*Bc*TUB). Data represent mean ± SD of three independent experiments. **d** Five-week-old Arabidopsis plant leaves were treated with 100 µM cellobiose, 100 µM AA9_COS or mock and callose was determined by aniline-blue staining 6 h and 24 h after treatments. The values of quantification relative to five leaves from five individual plants (n = 5) for each treatment and time point were analyzed. This experiment was repeated three times. Callose deposition was revealed by using ImageJ software and representative pictures are reported in **e**. f For each treatment the luminol-based H_2_O_2_ detection was performed in fourteen-day-old Arabidopsis seedlings (n = 10) from 0 to 40 min after AA9_COS and cellobiose treatments. Values are shown as mean ± SD. A representative experiment is shown in the figure. All the experiments were performed in three independent experiments with similar results. Lower case letters denote significant (p < 0.05) differences according to one-way ANOVA test with Tukey’s post-hoc multiple comparisons.

Callose deposition is a typical response that plants deploy to reinforce the plant cell wall during pathogenic attack^27^. To assess its deposition, leaves from five-week-old Arabidopsis plants were stained with aniline blue 6 h and 24 h following 100 µM AA9_COS and 100 µM cellobiose treatment (Fig. 2e). A substantial increase in callose accumulation was detected following AA9_COS treatment at both time points (Fig. 2d), while no callose spots were detected after cellobiose treatment (Fig. 2d, e, Supplementary Fig. 5a) ^33^. Interestingly, the dynamics and amplitudes in callose deposition due to AA9_COS treatment were proportional to time, acting stronger at 24 h than at 6 h (Fig. 2d, Supplementary Fig. 5a).

The transient and rapid generation of reactive oxygen species (ROS) is a hallmark of the response to MAMPs and chemical elicitor treatments^26,39,40^. Hence, the short-term (0-40 min) H_2_O_2_ generation was evaluated by the luminol-based assay (Fig. 2f). Interestingly, no measurable short- and long-term H O and O ^-^production was detected following AA9_COS treatment, indicating that AA9_COS triggers defense responses in a ROS-independent manner.

### Transcriptomic analysis of AA9_COS induced reprogramming in Arabidopsis

To gain insights into the dynamics of transcriptome reprogramming triggered by AA9-COS, Arabidopsis plants (fourteen-day-old) were treated with *i*) *Tt*AA9E products mixture (AA9_COS), and compared with treatments using *ii*) cellobiose, the only characterized cellulose derived product eliciting plants defense; the experimental set was completed with *iii*) mock treatments. To detect early responses^41^, the genome-wide transcriptomic analysis was performed 1 h after treatments using Arabidopsis 1.0 ST microarray chip (Affymetrix). Exogenous application of AA9_COS had a strong impact on the plant transcriptome landscape leading to changes in the expression of 545 genes (log_2_ Fold Change > 1 or < -1), of which 482 genes were upregulated, and 73 genes were downregulated compared to mock-treated plants (Fig. 3a). Conversely, cellobiose treatment modulated 352 genes, of which 283 were upregulated and 69 were down-regulated compared to mock-treated plants (Fig. 3a). A principal component (PC) analysis explained 53% of the combined variance and clearly separate mock, cellobiose and AA9_COS–treated samples (Fig. 3b). To assess the microarray data reproducibility, quantitative real-time PCR (qRT-PCR) analysis was performed on ten different genes (Supplementary Fig. 6). Microarray and qRT-PCR data showed a high and significant correlation (R^2^ = 0.89, FDR = 3.9e^-8^; Supplementary Fig. 6). To identify the biological processes in which the AA9_COS-responsive genes were involved, Gene Ontology (GO) and pathway enrichment analyses were performed (Supplementary Data 1). Most of the detected GO terms were related to plant defense, including responses to bacterial and fungal attacks (Fig. 3c). Among the enriched categories in AA9_COS-treated plants, there are *Innate Immune regulation, Cell wall modulation, Signaling, Response to hormones and Cell death regulation* (Fig. 3c). On the contrary, plants treated with cellobiose had the greatest enriched cluster associated solely with *Cell wall reorganization* (Fig. 3d). In summary, the enrichment analysis highlighted that the exposure of Arabidopsis to AA9_COS led to more complex and differential plant defense responses as compared to the cellobiose treatment.

**Fig. 3:**
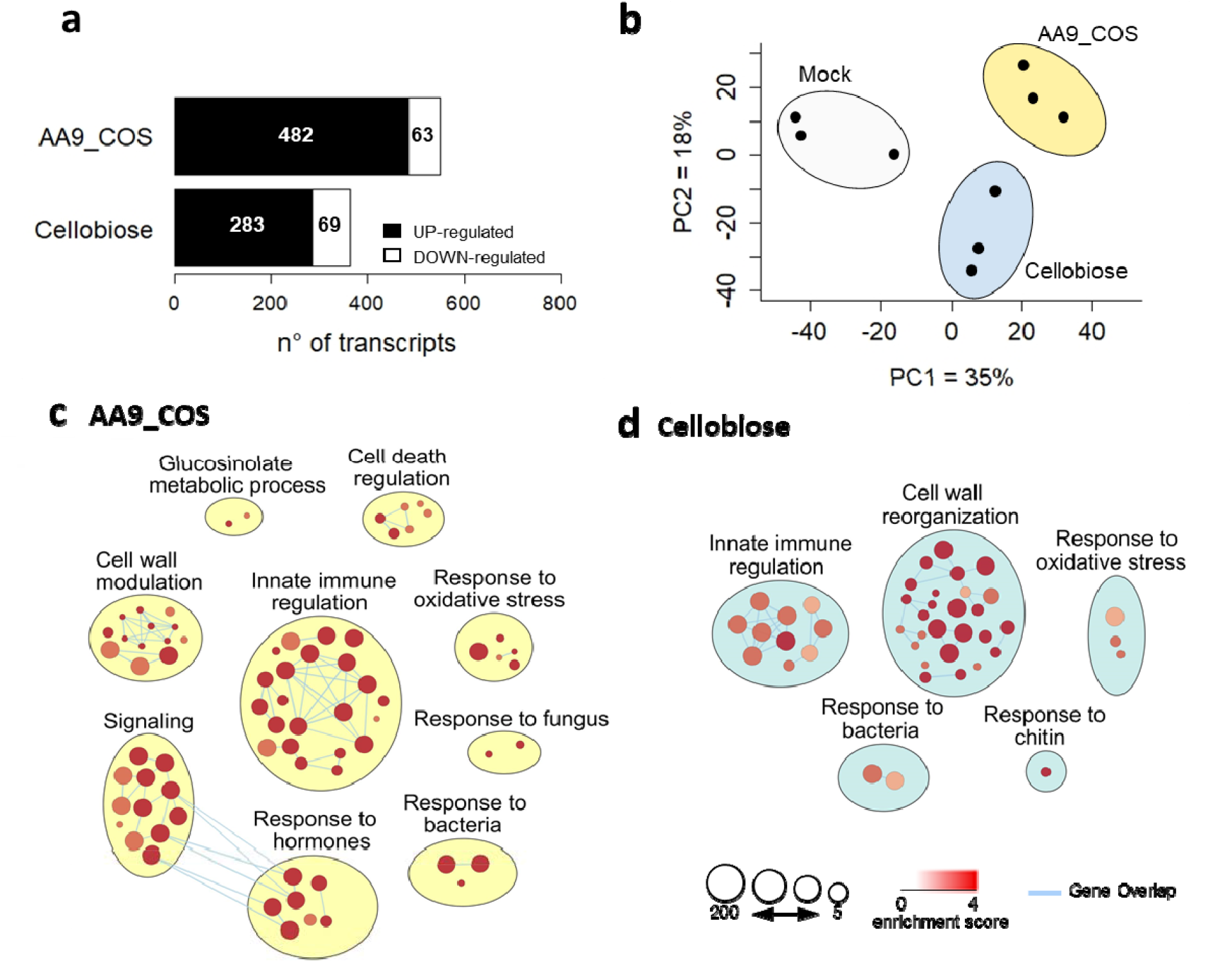
Global analysis of early transcriptomic changes in Arabidopsis seedlings treated with AA9_COS and cellobiose. **a** Fourteen-day-old Arabidopsis seedlings were treated with 100 µM AA9_COS, 100 µM cellobiose or mock; 1 h after treatment, fifteen seedlings were pooled and flash-frozen in liquid nitrogen. Three independent pools (n=3) for each treatment were produced and their relative transcriptome analysed. Genes featuring a log_2_ FC ≥ 1 or ≤ -1 compared to mock (p ≤ 0.05) were considered as differentially expressed. **b** Principal component analysis of transcriptome responses induced by AA9_COS (pale yellow), cellobiose (pale blue) and mock treatment (grey). The overall gene expression similarities between samples are visualized using two principal components (PC1 and PC2), representing 35% and 18% of the total variation, respectively. **c, d** Gene ontology (GO) enrichment analysis related to biological processes of 545 up-regulated by AA9_COS (c) and 352 genes up-regulated by cellobiose treatments (**d**). The GO analysis was performed by using g:Profiler software (Benjamin-Hochberg FDR < 0.05) and the EnrichmentMap function of Cytoscape^42^. Each node represents a pathway, edges represent overlapped genes between nodes and circle size represents the number of genes encompassed.

### LPMO-generated COS trigger powerful plant defense responses along with SA, JA, ET hormones production

To better understand the signaling events specifically induced by LPMO-derived products, we performed a hierarchical cluster analysis using AA9_COS and cellobiose transcriptome data. Clusters I and II were more responsive to AA9_COS than to cellobiose, whereas cluster III showed comparable gene expression between treatments (Fig. 4a, Supplementary Data 2). These clusters contained most of the core defensive marker genes like *FRK1, WRKY33* and *NAC DOMAIN CONTAINING PROTEIN 55 (NAC055)*^38^, as well as several genes encoding LRR-RLKs and Wall Associated Receptor Kinases (WAKs) (Fig. 4b and 4c, Supplementary Data 2). Cluster IV contained a few genes showing enhanced expression in cellobiose-treated than in AA9_COS-treated plants (Fig. 4a). These genes are associated with carbohydrate transport (Supplementary Data 2). Regarding the genes modulated by both AA9_COS and cellobiose and treatments, we observed that 26% (158 genes) of the total upregulated genes were common between treatments, whereas the percentage decreased to 15.7% (18 genes) for the shared downregulated genes (Fig. 4d). Interestingly, the -log_2_(FDR) analysis showed that AA9_COS-induced genes were more enriched for SA- and JA-responsive genes as compared to the cellobiose-induced genes, whereas a slight enrichment of ET-responsive genes was detected (Fig. 4e). These predictions were further proved by LC-MS quantification of SA, JA and gas-laser detection for ET, relative to AA9_COS-treated plants (Supplementary Fig. 7). About four-and eight-fold increase in SA and JA levels, respectively, were detected 24 hours after AA9_COS treatment as compared to mock. Instead, the ethylene was monitored in a time-course analysis from the immediate applications of AA9_COS until the following 24 h. An early two-fold increase of ET was observed 2 h after AA9_COS treatment as compared to mock and significantly higher ET level lasted till 4 h after treatment, which is typical for this volatile hormone^27^ (Supplementary Fig. 7).

**Fig. 4:**
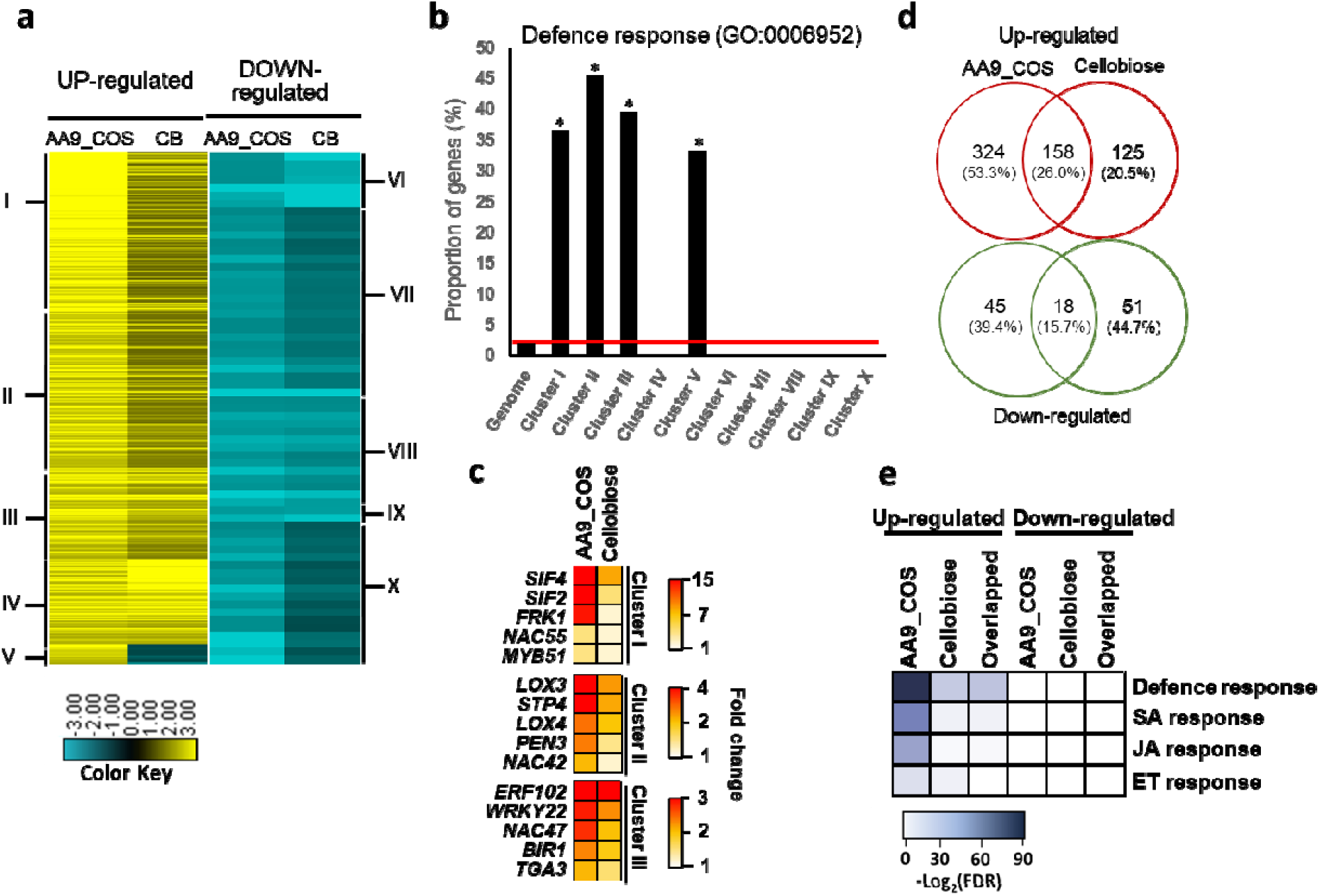
transcriptomic changes comparison between AA9_COS and cellobiose (CB) treatments. **a** Hierarchical clustering analysis of gene expression patterns that show significant expression changes as compared to mock (log_2_ FC ≥ 1 or ≤ -1, p ≤ 0.05) 1 h after 100 µM AA9_COS or 100 µM cellobiose treatments in fourteen-day-old Arabidopsis Col-0 plants. The clusters were divided into up-(yellow) or down-regulated (blue) genes. **b** The proportion of genes associated with the defense response GO-term (GO:0006952 Supplementary Data 3) was evaluated per cluster. **c** Heatmap of selected defense genes belonging to clusters I, II and III. d Venn diagram shows specific and shared up- and down-regulated genes induced by AA9_COS or cellobiose treatments. e The GO enrichment analyses performed with PSGEA indicated that defense- and phytohormones-related genes were upregulated both by AA9_COS and cellobiose treatments (Supplementary Data 4). The color key represents the enrichment significance shown as -log_2_ (FDR).

### AA9_COS modulates biotic-associated transcription factors

The identification of transcription factors TFs and their network is an important analysis for evaluating defense responses in expressomics data. Therefore, we performed a correlation network analysis (Pearson correlation > 0.9) using expression data with DEGs induced by AA9_COS treatment (log_2_ FC ≥ 1; Fig. 5). The analysis showed that WRKYs were the most represented TFs, followed by MYB, NAC and ERF families (Supplementary Table 1). *WRKY* genes known to play a role in PTI were found as central hubs in the network (*WRKY48, 47, 11, 72, 18, 29, 30* and *33*), together with *WRKY22* and *25* that were not previously associated with biotic stress^38^ (Fig. 5, Supplementary Table 1).

**Fig. 5:**
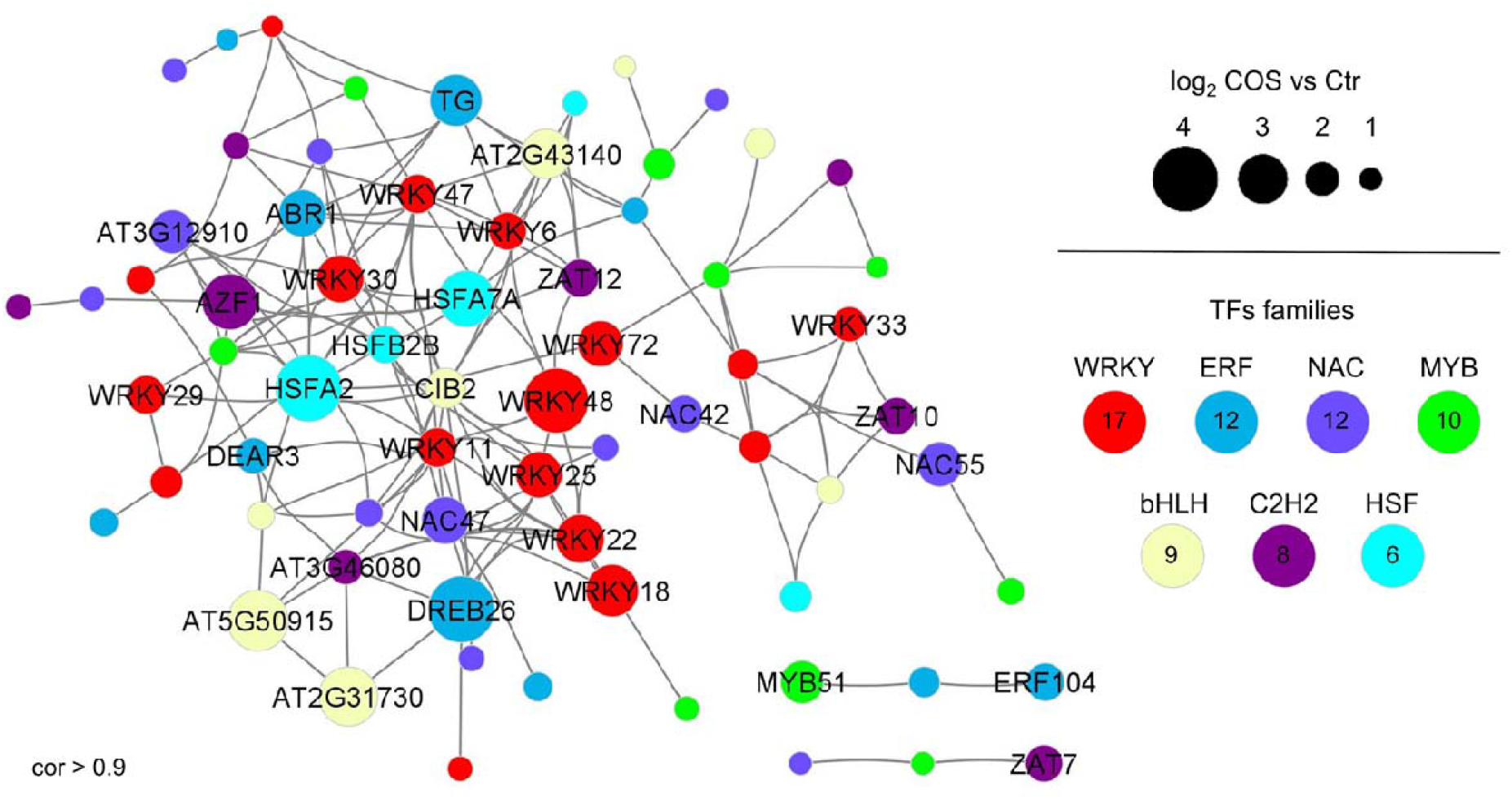
Correlation network of AA9_COS-upregulated transcription factors (TF). The correlation network was performed using significantly (p < 0.05) upregulated genes belonging to TF families. Grey edges indicate a positive correlation among TF genes whereas the circle sizes represent the different magnitude of gene expression reported as log_2_ FC AA9_COS vs mock. Only co-expressed upregulated TFs with r > 0.9 and TF families with at least five members up-regulated in AA9_COS were visualized in the network. The correlation network related to downregulated TFs is reported in Supplementary Fig. 8.

Moreover, AA9_COS induced the expression of several *NAC* genes (such as *NAC47 and 55*) (Fig. 5). In addition, members of the MYB TFs family were either up- or down-regulated by AA9_COS treatment (Fig. 5; Supplementary Fig. 8). Among the MYBs up-regulated by AA9_COS, we found *MYB51* gene, a major regulator of the MAMPs-dependent Indole-Glucosinolate accumulation in Arabidopsis^43^. Besides the canonical defense-associated TFs we also observed members of bHLH, C2H2 and HSF families being modulated by AA9_COS (Fig. 5). Interestingly this analysis provided that several marker genes are specifically enriched upon cello-oligosaccharide treatment probably indicating more suitable markers to evaluate native and oxidized COS immunity induction.

### Carbohydrate transport proteins are specifically modulated by AA9_COS

Other classes worthy to be reported are those of sugar transporters such as Sugar Transport Proteins (STPs), SWEET, and Sucrose Transport Proteins (SUCs), well-known to be hijacked by pathogens to favour sugar efflux from the cytosol to the apoplasm^44,45^. Several genes encoding sugar importer proteins such as STP4, -13, -1, were induced by AA9_COS mixture. On the contrary, genes encoding the most important SWEET bi-directional transporters *(SWEET 2, 16, 17)*, involved in sugar export from the cytoplasm to the apoplast, were downregulated by treatment (Supplementary Table 2). These data suggest that AA9_COS might also counteract the pathogen invasion by leading plants to sequestrate edible resources for phytopathogens.

### *STRESS INDUCED FACTOR 2 and 4* and other LRR-RLKs regulate the AA9_COS-induced defense gene expression

To further decipher the molecular mechanism underlying defence activation by AA9_COS, the transcriptomic data were screened for Pattern Recognition Receptors (PRRs) and Receptor-Like Kinases (RLKs) proteins, pivotal players in DAMPs signaling^21^ (Supplementary Table 2). Two genes encoding Leucine-Rich Repeat RLKs (LRR-RLKs), namely *STRESS INDUCED FACTOR 2* and *4 (SIF2, SIF4)*, were strongly upregulated by AA9_COS and represented in cluster I (Fig. 4c). In addition, two other RLKs, such as *BRI1-ASSOCIATED RECEPTOR KINASE 1 (BAK1)* and *THESEUS 1 (THE1)*, key players for flagellin perception^22^ and cell wall integrity sensing^4^, respectively, were triggered by AA9_COS (Supplementary Table 2) and selected for further studies. To evaluate the potential role played by these LRR-RLKs in signaling the AA9_COS-dependent defence gene expression, the *sif2, sif4, bak1* and *the1* KO mutants were treated with 100 µM AA9_COS. The expression levels of three defense marker genes, *FRK1, WRKY22* and *33*, along with *SIF2* and *SIF4* genes, were evaluated 1h after treatment (Fig. 6). Interestingly, in all the AA9_COS treated mutant lines, the expression of *FRK1* and *WRKY22* was equal to mock-treated Col-0 plants (Fig.6). A slight but not significant increase in *WRKY33* expression was detected in AA9_COS-treated *sif2, bak1* and *the1* KO lines as compared to mock-treated plants. However, the level of expression of these genes in the mutant background were significantly lower as compared to what observed in AA9_COS-treated Col-0 plants (Fig. 6). Altogether, these data suggest that these *BAK1* and *THE1* plasma membrane localized RLKs might interact singularly or in combination with *SIF2* and/or *SIF4* to create an active complex for the perception of AA9_COS.

**Fig. 6:**
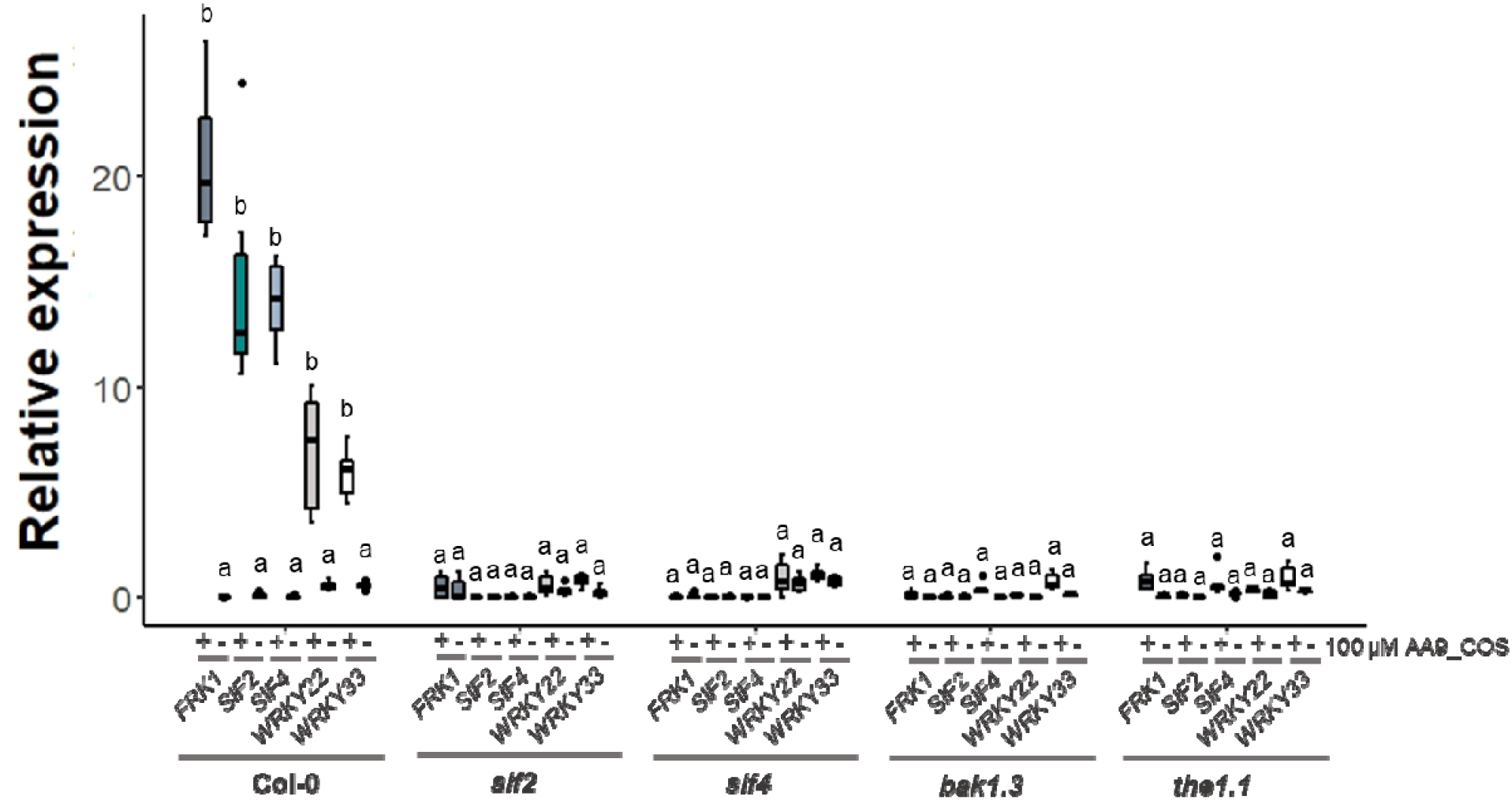
Involvement of selected RLK receptors in signaling the AA9_COS-dependent defense gene expression. The expression of defense marker genes was evaluated in *sif2, sif4, bak1*.*3* and *the1*.*1* loss-of-function Arabidopsis seedlings. Fourteen-day-old seedlings were treated with or without 100 µM AA9_COS and qRT-PCR was carried out 1h after treatment. Median values are plotted in the boxes with data generated from three independent pools (n = 3) of fifteen seedlings each. One-way ANOVA test was performed; lower case letters denote significant (p < 0.05) difference between mock and AA9_COS treated plants.

### *STRESS INDUCED FACTOR 2* and *4* regulate the AA9_COS-induced camalexin accumulation independently to MPK3/6 phosphorylation

The tryptophan-derived camalexin is a well-known antimicrobial metabolite produced by plants during pathogenic infection, i.e. *B. cinerea*^29,46,47^. Camalexin was quantified in fourteen-day-old Col-0, *sif2* and *sif4* KO mutants 24 hours after 100 µM AA9_COS, 100 µM cellobiose or mock treatment. The two latter yielded comparable levels of camalexin (mock = 2.4 ± 1.4 ng/g, cellobiose = 3.7 ± 1.8 ng/g), whereas AA9-COS yielded 35.0 ± 5.4 ng/g in Col-0 plants. On the contrary, no camalexin production, along with an increase of *B. cinerea in planta* growth, was observed in both sif2 and sif4 AA9_COS-treated plants (Fig. 7a, Supplementary Fig. 9). Given the involvement of the *MPK3-MPK6-WRKY33* module in the activation of the camalexin biosynthetic pathway ^48^, the expression of these genes was measured in Arabidopsis Col-0 plants 1 h and 24 h after treatment with 100 µM AA9_COS or 100 µM cellobiose. Generally, treatments with AA9_COS yielded stronger inductions: 1h after treatment a significant (p < 0.05) induction of *MPK3/6, WRKY33, PAD3* and *PEN3* was detected, while at 24 h after treatment only *MPK3* gene was maintained expressed (Fig. 7b relative to fourteen-day-old plants). Instead for cellobiose treatment a lower induction was observed: except for *MPK3/6*, at 1h after treatment a slight expression was observed for *WRKY33, PAD3* and *PEN3* genes as compared to mock-treated plant, whereas no significant (p > 0.05) differences were detected at 24 h (Fig. 7b). Finally, similar trends of *MPK3*/6 and *WRKY33* gene expression were observed when comparing two-week-old with five-week-old plants (Supplementary Fig. 10). Moreover, the MPK3/6 gene expression activation was completed by immunoblotting analysis. Early MPK3/6 phosphorylation was detected after 100 µM AA9_COS treatment, whereas no phosphorylation was observed upon treatments with 100 µM cellobiose in Col-0 plants (Fig. 7c). Therefore, the MPK3/6 proteins and their phosphorylated forms (pMPK3/6) were hybridized in Col-0, *sif2* and *sif4* mutant lines. After AA9_COS treatment, we observed a signal for the pMPK3/6 both in Col-0, *sif2* and *sif4* KO lines, even if a slight decrease in pMPK3/6 forms was observed in sif4 KO lines (Fig. 7c). Altogether these data indicate the SIF2/4 LRR-RKs as keys regulators to signal the AA9_COS-dependent response activation.

**Fig. 7:**
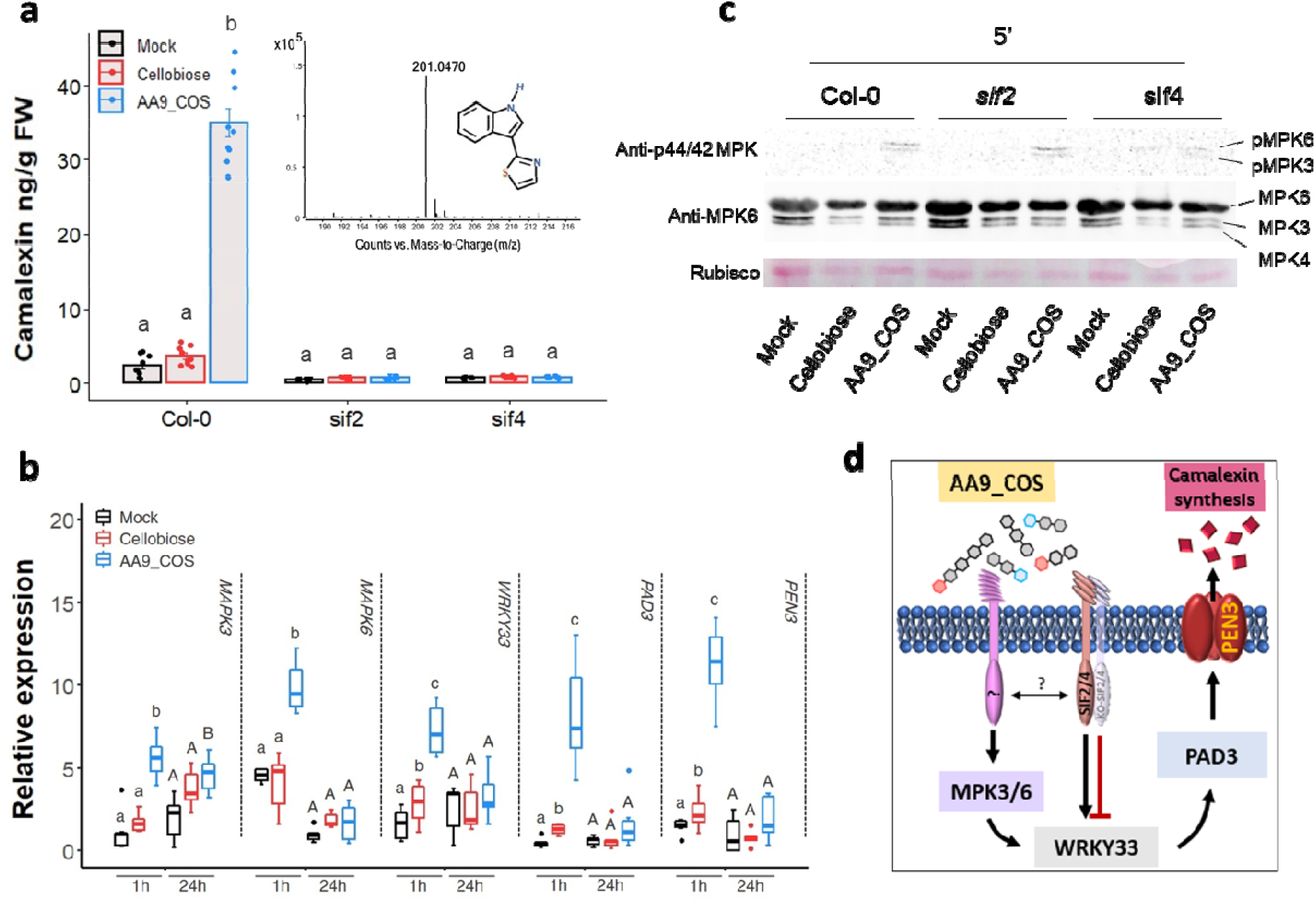
AA9_COS induces camalexin biosynthesis and requires *SIF2* and *SIF4* LRR-RLK genes. **a** Camalexin was quantified in fourteen-day old Col-0, *sif2* and *sif4* Arabidopsis seedlings by LC-MS 24 h after 100 µM AA9_COS and 100 µM cellobiose treatments. Bar plots represent the mean ± SD of data generated from three independent experiments (n = 3). One-way ANOVA with Tukey’s post-hoc multiple comparisons (p ⍰<⍰ 0.05) were performed; lower case letters denote differences. The mass of camalexin is reported. **b** The expression of genes involved in camalexin signaling and biosynthesis was evaluated in fourteen-day old Col-0 Arabidopsis seedlings 1 h and 24 h after 100 µM AA9_COS, 100 µM cellobiose and mock. Median values are plotted in the boxes with data generated from three independent pools (n = 3). Each pool consisted of fifteen seedlings. One-way ANOVA with Tukey’s post-hoc multiple comparisons (p < 0.05) was performed; lower case and capital letters denote difference at 1 h and 24 h following treatments, respectively. **c** Immunoblotting analysis showing the early activation of MPK3 and MPK6 protein kinases after indicated treatments. Fourteen-day old Col-0, *sif2* and *sif4* Arabidopsis seedling were treated with 100 µM AA9_COS, 100 µM cellobiose and mock and samples were collected five minutes after treatment. Western blot analysis was carried out using the following antibodies: anti-p44/42 MPKs antibody (Cell Signaling Technology, active forms), anti-MPK3 (Sigma, total proteins) and anti-MPK6 (Sigma, total proteins). Loading proteins were detected using the Ponceau S staining and Rubisco is shown in the bottom panel. Three independent experiments were performed with similar results. **d** Proposed model for the AA9_COS-regulation of camalexin biosynthesis and signaling.

## Discussion

In this study, we found that the mixtures of oxidized and native cello-oligosaccharides produced by the fungal cellulose oxidizing LPMO AA9 enzyme (AA9_COS) are eliciting plant immune responses in higher and complete extend than the solely cellobiose, a native cello-oligosaccharide released instead by glucosyl hydrolases enzymes (hydrolytic CWDEs). The LPMO-driven oxidation of cellulose generates a mixture of C1-, C4-oxidized and native COS (Fig. 1c) which leads to a wide transcriptome reprogramming and activation of immune responses in Arabidopsis (Fig. 3, 5 and 7). We observed that the AA9_COS-dependent defense response activation conferred increased resistance to the necrotrophic phytopathogens B. cinerea both in Arabidopsis and tomato plants (Fig. 2, Supplementary Fig. 3). Additionally, we discovered that two plant LRR-RLKs, namely SIF2 and SIF4 are required to fully signal the AA9_COS-dependent responses (Fig. 6), pointing toward their possible role in sensing biotic or abiotic stresses or alternatively, DTI and CWI maintenance mechanisms. As indicated previously, DTI activation and CWI sensing are two distinct mechanisms implicated in monitoring chemical and mechanical alterations of the cell wall. Nevertheless, the partially overlapped pathways controlling these two mechanisms make it hard to distinguish one from another^3–5^. In plant-pathogen interactions, the role of DAMPs resulting from the activity of hydrolytic enzymes on pectin and hemicelluloses is well established^30,31^. Classic examples are the well-characterized pectin-derived OGs and xylans from hemicelluloses^31,32,41,49^. Conversely, little is known about the signaling pathways and defence responses triggered by perturbations of the main load-bearing component of the cell wall, i.e., cellulose.

Fungal LPMOs possess the unique feature to not only produce diffusible DAMP signals in the form of peculiar mixtures of native, C1- and/or C4-oxidized oligosaccharides, but also to weaken the crystalline regions of the cell wall, causing loss of structural integrity^8,50^. Indeed, LPMOs have been recently proposed to act earlier than other CWDEs during cellulose degradation^18^, hence in light of the results here presented, it might play a pivotal role in plant-pathogen interactions. Previously, it was revealed that during pathogenicity on different hosts, the necrotrophic fungus *B. cinerea* deployed three LPMOs, all from the AA9 subfamily^7^. In this work, we studied the transcription pattern of the entire BcAA9 gene family during the infection of Arabidopsis plants by *B. cinerea* (Supplementary Fig. 1). The sequence alignment of the *Bc*AA9 catalytic domains against structurally resolved fungal AA9s enzymes showed that the majority of expressed proteins potentially have C1/C4 oxidative activity (Fig. 1a). Therefore, the *TtAA9E* from the saprophytic fungus *T. terrestris* was chosen from our collection to produce a representative AA9_COS mixture (containing either native and C1 and/or C4 oxidized cello-oligosaccharides) as predicted by the phylogenetic tree analysis related to *BcAA9* genes (Fig. 1a, b, Supplementary Fig. 1). The mixture so produced (AA9_COS) contained native and oxidized (aldonic acids and gemdiols) oligosaccharides of various degrees of polymerization (DP). Indeed, the LPMO-derived products, rather than the single native oligosaccharide, better represent the blend of COS released from the plant cell wall polysaccharides naturally occurring in a pathogenic attack. Therefore, the study of this more complex blends of COS provides a valuable biological output that was further characterized in the rest of the work.

In the graphical model (Fig. 8) we summarized all the major results here obtained from the transcriptomic reprogramming, cellular and physiological changes that conferred Arabidopsis higher resistance to the pathogenic fungus *B. cinerea* upon AA9_COS treatments. Several PRRs and RLKs were modulated by AA9_COS treatment (Supplementary Table 2) and the genes encoding two LRR-RLKs, *SIF2* and *SIF4*, were highly expressed (Fig. 4c, Supplementary. Table 2). SIF2 interacts with the FLS2-BAK1 PRR complex and regulates either MAMPs-dependent PTI pathway, involving MAPK kinases, *WRKY* TFs and *FRK1* and pathogen-dependent stomata closure^51,52^. Moreover, through a meta-data analysis, we recently identified *SIF4* as constantly expressed during several combined biotic and abiotic stress conditions^53^. In the results here presented, we showed that both *SIF2 and SIF4* play a crucial role in signaling the AA9_COS-dependent responses such as camalexin production. Moreover, necrotrophic growth of *B. cinerea* was significantly higher in *sif2* and *sif4* KO mutants if compared to Col-0 plants probably due to a lesser amount of camalexin being produced in the mutants. Moreover, treatments with AA9_COS could still aid the plants against the pathogen in retarding its growth although only partially (Supplementary Fig. 9). Indeed, the MAPKs gene expression and phosphorylation were rapidly triggered by AA9_COS treatment (Fig. 7b, c), which is in agreement with what was previously observed following OGs and native COS treatments^54^ triggering camalexin^55^ production and enhancing resistance to the fungus *B. cinerea*^37^. The production of the antimicrobial compound camalexin is therefore the final output of defense mechanisms deployed by plants upon metabolic reprogramming involving several anabolic enzymes^27^. Our results showed the induction of key regulatory genes as well as the actual synthesis of this metabolite in AA9_COS-treated plants (Fig. 7b, d). Moreover, a vast *de-novo* deposition of callose conferring resistance to *B. cinerea* (Fig. 2a-e) was observed. In contrast, these basal defenses were not triggered by cellobiose alone (Fig. 2a-e) ^33^. Interestingly, no expression of three PTI-marker genes, as well as a null accumulation of camalexin, was detected in both *sif2* and *sif4* KO lines after AA9_COS treatment (Fig. 6 and 7a), indicating these LRR-RLKs as potential key players for AA9_COS-regulated defense induction^51^.

**Fig. 8:**
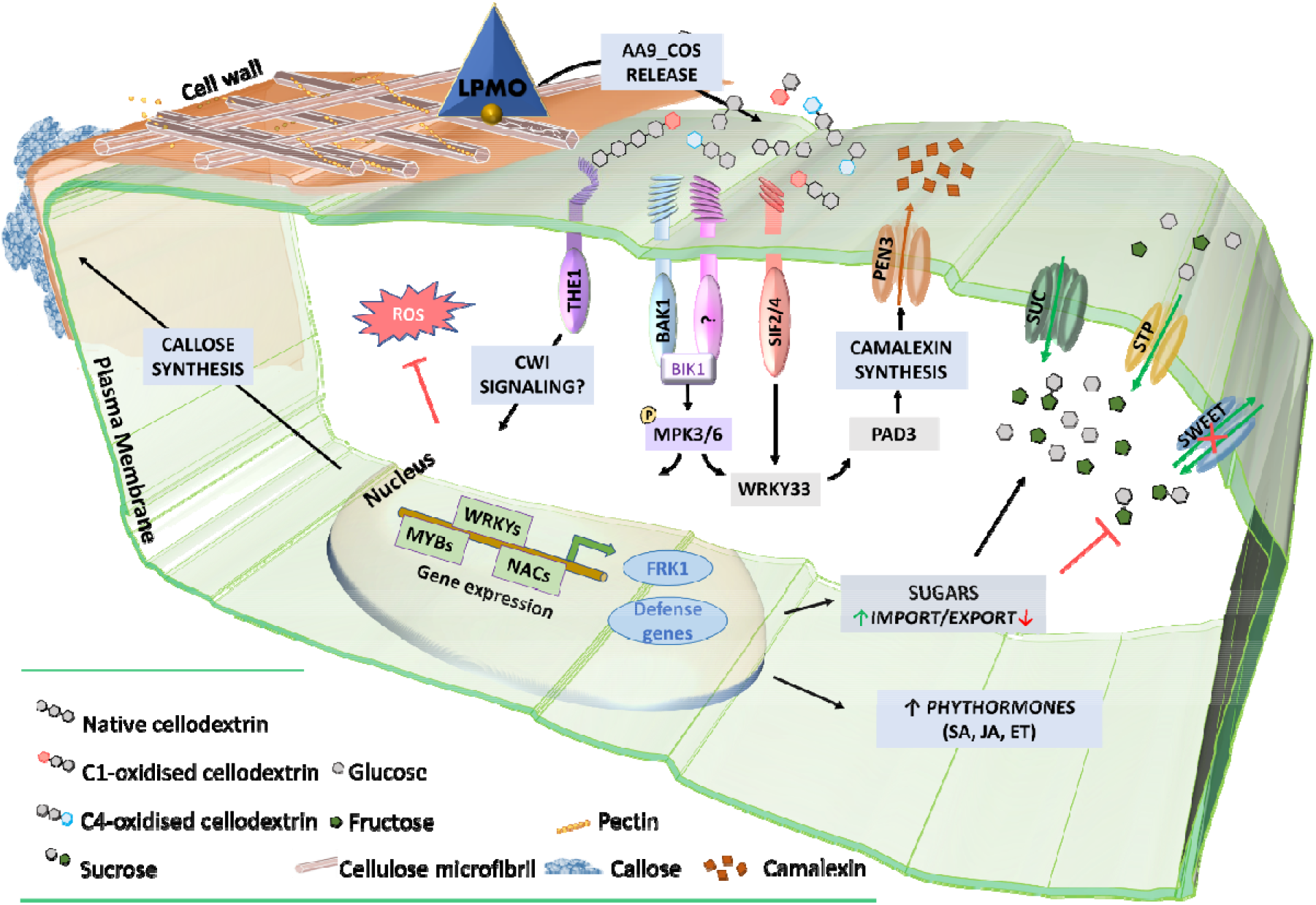
schematic model for AA9_COS perception, signaling and defense responses in Arabidopsis. The degradation of plant cell wall polysaccharides by microbial LPMOs leads to the release of native and oxidized cello-oligosaccharides such as AA9_COS. In the apoplast, plasma membrane-anchored PRR proteins, i.e., SIF2/4, BAK1 and THE1, after DAMPs sensing activate ROS-independent DTI responses which include callose deposition and phytohormones accumulation. Consequently, the recognition of AA9_COS induces a deep transcriptional reprogramming, including TFs belonging to the WRKY, MYB and NAC families, as well as the phosphorylation of MPK3 and MPK6. This, in turn, leads to a remarkable synthesis of camalexin. Additionally, genes encoding for sugar transporters are modulated by AA9_COS treatment, with the induction of cytoplasmic sugars import (SUC and STP) and the inhibition of SWEET transporters involved in sugar export from the cytoplasm to the apoplast.

Furthermore, only a slight decrease in MPK3/6 phosphorylation was observed in *sif4* mutants upon AA9_COS treatment as compared to wild type plant (Fig. 7c). This is in accordance with previous data showing that in *sif2* mutants, the MAPK3/6 cascade is not impaired upon *Pst* DC3000 infection^51^, suggesting that these two RLKs might regulate basal defenses in an MPK3/6-independent manner. Interestingly, a null AA9_COS-dependent *FRK1, WRKY22* and WRKY33 gene expression was observed in *bak1* and *the1* loss-of-function mutants, impaired for the membrane-receptors required for flagellin^22^ and CWI sensing^3,4^, respectively (Fig. 6). These results suggest that AA9_COS mixture might play a role also in signaling mechano-derived damages of cell wall^58,59^, like those caused by abiotic stress, besides canonical immune responses.

We hypothesized that the presence of native and oxidized oligomers in the AA9_COS, leading to a stronger response than cellobiose although equal in molar number, is probably due to a synergistic effect and simultaneous perception by various membrane receptors (Fig. 3b-d; Fig. 4d). Several genes with a major role in plant defence, i.e. JA- and SA-regulated genes as well as their biosynthetic pathways, were positively modulated by AA9_COS treatment (Fig. 3b-c, e). This was also confirmed by the higher amount of SA, JA quantified after 24 hours from AA9_COS treatments as compared to mock (Supplementary Fig. 7). The production if the volatile hormone ethylene was instead detected shortly (1-2 hours) after AA9_COS treatment and its emanation dissipated 4 hours later (Supplementary Fig. 7). Phytohormones such as SA, JA and ET are key chemical players governing the PTI and DTI activation^27^. Indeed, earlier investigation based on carbohydrates elicitors such as OGs and β-1,3-glucan, have been reported to regulate plant defenses via SA- and JA-dependent expression and production^41,49^. On the other hand, ET is generally associated to PTI response. For example an identical kinetics of ET production starting from 1 h and peaks at 4 h was observed Arabidopsis seedlings treated with Flg22^27^. Altogether these data confirm that AA9_COS might trigger DTI and PTI responses in Arabibopsis.

Finally, sugar transporters are also impacted by AA9_COS treatment. The transcriptome analysis showed that AA9_COS positively modulated sugar importers of the classes SUC and STP, whereas SWEETs efflux carriers, involved in exporting hexoses from the cytosol to the apoplast, were inhibited (Fig. 8, Supplementary Table 2). Therefore, it seems that after perceiving AA9_COS plant cells may retain sugars intracellularly as a strategy to counteract pathogen attacks^44^. On the other hand, it has been previously hypothesized that plants regulate the growth-immunity trade-off by using a homeostatic mechanism governed by the Berberine Bridge Enzyme (BBE) family^54^. More specifically, it has been reported that the oxidation of cellooligosaccharides, such as cellotriose and cellotetraose (DP3-4), by BBE-CELLOX when overexpressed in Arabidopsis, inactivate the elicitor properties of these native COS^54^. Intrigued by this observation, we treated Arabidopsis plants with cellotrionic acid (DP3) and the shorter cellobionic acid (DP2) produced *in vitro* by using the enzyme cellobiose dehydrogenases (CDH). We observed that the C1-oxidized cellobionic acid could induce an higher expression of *MPK3, RBOHD, CYB81F2, LOX1 and WRKY33* genes as compared to control plants, while in agreement with Locci and co-workers^54^, a null expression of *WRKY30, CYB81F2, RBOHD and LOX1* genes was measured following cellotrionic acid treatment (Supplementary Fig. 11). Our data support the fact that oxidized COS maintains the defense elicitor properties also when applied singularly without their native counterpart and pointing towards the shorter cellobionic acid (DP2) as the most active form in eliciting plant immunity among the DP2-4.

Oligosaccharides and products of cell wall depolymerization are attracting a great deal of interest in agrobiotechnology for their ability to elicit plant immune defenses in an environmental and human friendly mechanism. Till now, the only known COS eliciting plant immunity were those derived by hydrolytic enzymes (CDWE), mostly active on hemicellulose and pectin such as polygalacturonase and xylanase, and only recently cellulose was considered. To the best of our knowledge, we are reporting a seminal study showing that the new redox enzyme LPMO of the AA9 family are also able at triggering plant immunity. The mixture of cellooligosaccharide released by the carbohydrate oxidative enzymes largely known as AA, although in large presence in phytopathogens genomes and proteomes, were not investigated as plant defense elicitors yet. Given the enormous variety of LPMO active on several plant polysaccharides, i.e., cellulose, starch, hemicellulose, xylans, xyloglucans and β-glucans, we believe that their derived blends of native and oxidized oligosaccharides (virtually myriad combinations) might represent a fertile source of new bio-derived molecules to be used for plant protection or defenses priming against various pathogens.

## Methods

### *Tt*AA9E cloning, expression and oligosaccharide characterization

The synthetic gene encoding *Tt*AA9E gene from *Thermothielavioides terrestris* (ACE10234.1), obtained from Eurofins Genomics (Ebersberg, Germany), was cloned into a pPICZα-A modified expression vector, as previously described^16,36,58^. The resulting plasmid (∼10 μg of pPICZT::lpmo) was inserted by transformation into *P. pastoris* X-33 and the secreted proteins were purified by affinity chromatography (HiTrap Chelating HP column) (GE Healthcare). Eluted proteins were copper saturated with Cu(II)SO4 and applied to a 120 mL Superdex 75/16 column (GE Healthcare) to remove the excess of copper. Finally, proteins were concentrated by ultrafiltration. The total protein was quantified using the Bradford method and purity was analyzed by SDS-PAGE. The cellulosic substrate PASC (0.5 % w:v), obtained as previously described^13^, was incubated with 1 μM *Tt*AA9E in 10 mM sodium acetate buffer (pH 5.0) using 500 µM ascorbic acid as a reducing agent. The reaction was incubated in agitation at 800 rpm at 50 °C for 72 h. Five reaction tubes were prepared each time individually analyzed via HPEAC and later pooled and purified with molecular filter (3KDa cut-off, Vivaspin) stored at -20 °C until further used. HPAEC chromatography was done using a Dionex ICS6000 instrument equipped with pulsed amperometry detection (PAD) and a CarboPac PA1 column (2×250 mm). A large amount of AA9_COS was produced and available upon reasonable request.

### Preparation of oligosaccharides for plant treatments

The AA9_COS mixture obtained as described above was adjusted by adding Milli-Q water to 100 µmol and filtered through 3KDa molecular sieve before plant treatments.

### Plant Material, growth conditions and pathogen infections

*Arabidopsis thaliana* Columbia-0 (Col-0) wild type, *sif2* (SALK_209134C) and *sif4* (SALK_208927C) KO T-DNA insertional lines were obtained from the Nottingham Arabidopsis Stock Centre, *bak1* and *the1* loss-of-function line were kindly provided by Prof. Herman Höfte (IJPB, INRAE-Versailles, France). Seeds were surface sterilized by subsequent treatment with 70% (v:v) ethanol and 3% (v:v) bleach solution in agitation for 3 and 5 min, respectively. Seeds were then washed three times with distilled water and stratified for 3 days at 4 °C in the dark after sowing.

For the “*in planta”* growth test of *B. cinerea*, 5-week-old Col-0, *sif2, sif4* and two-month-old tomato plants were germinated in soil and grown in a controlled chamber (same growth conditions as above). Fifteen leaves from three independent plants were treated by applying 40 - 50 µL of filtered solutions per leaf. At the indicated timepoints, treated leaves were inoculated with 10-μL-droplet of *B. cinerea* spores (5·10^5^ spores mL^-1^), and three days after infection leaves were pooled, flash-frozen in liquid nitrogen and stored at – 80°C. Genomic DNA from ground-frozen samples was extracted and fungal biomass was quantified as previously described by qRT-PCR analysis using primers specific for *B. cinerea (BcTub)* and Arabidopsis *(AtSK11)* ^37^. Three independent pools (n = 3) were so replicated.

The *B. cinerea* symptomatology was performed in five detached leaves collected from three 5-w-old Arabidopsis or 2-month-old tomato plants (n = 3) by inoculating 10-μL droplet (5 10^5^ spores mL^-1^) per leaf. Three days after fungus inoculation, pictures were taken with a photo camera and the areas of the developing necrotic lesions were analysed by ImageJ software. This experiment was repeated three times.

The *B. cinerea* B05.10 strain was grown on potato dextrose agar medium at 21 °C, with a photoperiod of 12 h light (100 µmol photons m^-2^ s^-1^)/12 h darkness.

### Sequence alignment, phylogenetic analysis and structural model

The annotated AA9s from *Botrytis cinerea* BCIN_15g03140, BCIN_05g08230, BCIN_06g00480, BCIN_12g03920, BCIN_03g05890, BCIN_11g05360, BCIN_09g06750, BCIN_09g06730, BCIN_02g02040 and BCIN_06g07050 were aligned with the AA9s biochemically and structurally characterized *Af*AA9B (PDBid 5×6A), *Cv*AA9 (PDBid 5NLT), *Hj*AA9A (PDBid 5O2W), *Hi*AA9B (PDBid 2VTC), *Ls*AA9A (PDBid 5ACF), *Mt*AA9D (PDBid 5UFV), *Nc*AA9A (PDBid 5FOH), *Nc*AA9C (PDBid 4D7U), *Nc*AA9F (PDBid 4QI8), *Nc*AA9M (PDBid 4EIS), *Pc*AA9D (PDBid 4B5Q), *Ta*AA9A (PDBid 2YET) and *Tt*AA9E (PDBid 3EII) using MUSCLE^59^. Phylogenetic analysis was performed with MEGA software^60^ version 10.1.7 using the neighbour-joining method. The consensus tree was inferred using a bootstrap of 1,000 replicates. The structural model of BCIN_12g03920 was generated using the Swiss-Model Automated Comparative Protein Server and using LsAA9A (PBDid 5ACF) as template. Protein structures were visualized with PyMol Molecular Graphics Systems (Version 1.5.0.4 Schrödinger, LLC, New York, NY, USA).

### RNA extraction, cDNA preparation and qRT-PCR analysis

Total RNA was extracted from 100 mg of ground-frozen samples using the Spectrum™ Plant Total RNA Kit (Sigma-Aldrich) and the RNA purity was determined through NanoDrop 2000 UV-Vis Spectrophotometer (Thermo Scientific, Loughborough, UK). First-strand cDNA was synthesized by using SOLIScript RT cDNA synthesis MIC (Solis Biodyne) and qRT-PCRs were performed using SYBR Selected MasterMix 2x (HOT FIREPol^®^ SolisGreen^®^ qPCR Mix, Solis Biodyne), following the manufacturer’s protocol in a MyiQ real-time PCR detection system (Bio-rad). The cycling conditions consisted of an initial 7 min at 95 °C, followed by 40 two-step cycles at 95 °C for 5 s and 60 °C for 30 s. Melting curve analysis was performed after cycle completion to validate amplicon identity. Relative expression levels for plant gene expression were calculated following the standard curve-based method by using *ACTIN2* and *Elongation Factor-1α* as reference genes ^61^. Relative expression levels for *B. cinerea* AA9s gene in planta and in PDA medium growth were calculated following the standard curve-based method by using *BcTUB* as reference gene. Gene-specific primers used in this work are listed in Supplementary Table 3. All the qRT-PCR were measured twice from 3 independent (n=3) pools each consisting of 15 leaves.

### Microarray Hybridization and Data Analysis

RNA sample preparation for microarray hybridization was carried out as described in the Applied BiosystemsTM GeneChipTM Whole Transcript (WT) PLUS Reagent Kit User Guide (Thermo Fisher Scientific, Waltham, MA, USA). Briefly, 200 ng of total RNA was used to generate double-stranded cDNA. Twelve µg of subsequently synthesized cDNA was purified and reverse transcribed into single-stranded (ss)cDNA, where unnatural dUTP residues were incorporated. Purified and labeled sscDNAs were hybridized to Arabidopsis Gene 1.0 ST arrays (Affymetrix) and the fluorescent signals were measured with an Applied BiosystemsTM GeneChip Scanner 3000 7G System. Three biological replicas were hybridized for AA9_COS, cellobiose and mock-treated RNA samples extracted from three independent pools (n = 3) for each treatment consisting of fifteen seedlings per pool. Sample processing was performed at a Genomics Core Facility, “KFB - Center of Excellence for Fluorescent Bioanalytics” (Regensburg, Germany). Normalized probe set signals in the log_2_ scale were calculated by using the RMA algorithm (Applied Biosystems GeneChip Expression Console v1.4 software) and the Differentially Expressed Genes (DEGs) were selected using two criteria: log_2_ fold change ≥ 1- or ≤ -1 and Student’s t-test p-value ≤ 0.05. Average linkage hierarchical clusters of DEGs were generated by using Cluster 3.0 and visualized by Java Treeview.

### Gene ontology, pathway enrichment analysis and network analysis

Gene Ontology analysis related to the biological process was performed by Cytoscape software using upregulated genes (log_2_ fold change > 1, p <0.05) through g:Profiler software using Benjamini and Hochberg FDR (< 0.05) as filtering step accordingly to^62,63^. In addition, to better visualize the results of the GO analysis, terms were clustered by using the “Autoannotate” in Cytoscape tool, thus obtaining 10 clusters associated with defense responses. The network analysis was performed using the “Expression Correlation” app of Cytoscape software^64^.

### Callose staining

Callose deposits quantification was performed as previously described^39^. Briefly, for each treatment five leaves from five 5-w-old Arabidopsis plants were analyzed (n = 5). This experiment was performed three independent times. Leaves were stained using 0.01% (w:v) aniline blue in 150 mM K_2_HPO_4_ (pH 9.5) buffer for 30 min and then de-stained in a lactophenol clearance solution overnight. Leaves were examined by stereofluorescence microscopy with an Azio Zoom V.16 (Carl Zeiss Inc., Oberkochen, Germany) and callose spots were quantified using ImageJ software. All staining experiments were repeated three times and representative images were selected.

### Hydrogen peroxide detection assays

Hydrogen peroxide (H_2_O_2_) was assayed with Arabidopsis plant cultivated in vitro or in soil. The H O in situ detection was performed as previously described^65^. Briefly, for each treatment, ten leaves from three 5-weeks-old Arabidopsis plants were droplet-treated (40 – 50 µL per leaf) with indicated compounds. At 24 h after treatment leaves were gently vacuum-infiltrated (five min) with 1 mg mL^-1^ 3,31-diaminobenzidine (DAB) dissolved in 10 mM sodium phosphate buffer and 0.05 % (v:v) Tween 80. The staining reaction was terminated 5 h after DAB infiltration, and leaves were fixed in ethanol:glycerol:acetic acid 3:1:1. Chlorophyll was removed by several washing steps with 70% (v:v) ethanol and pictures were taken with a stereomicroscope (Discovery V8, Zeiss).

Elicitor-induced H_2_O_2_ was also detected by luminol-peroxidase-based. Briefly, twenty leaves from ten fourteen-day-old Col-0 seedlings were cut and incubated in ddH_2_0 overnight. Also discs from 5-weeks-old Arabidopsis were cut and assayed similarly. Time course luminescence was evaluated using The SpectraMax iD3 Multi-Mode Microplate Reader (Molecular devices) with an integration time per well of 1 s following application of indicated compounds.

### Superoxide detection assay

Superoxide anion (O_2_ ^-^) was assayed by performing nitro blue tetrazolium (NBT) assay^65^.

### LC-MS for camalexin quantification and hormones

For each treatment, camalexin was extracted from a pool of fifteen seedling of fourteen-day-old Arabidopsis, following method^37^.

Detection and quantification of JA and SA were performed at the Plant Observatory-Chemistry and Metabolomics platform (IJPB/INRA, Versailles, France), as previously described^39^.

### Protein extraction and immunoblotting

For MAPK3/6 immunoblotting, fifteen seedlings 14-days-old Col-0, *sif2* and *sif4* were treated with 100 µM AA9_COS, 100 µM cellobiose or mock. Leaves were collected 5 min after treatment, immediately frozen in liquid nitrogen and stored at -80 °C for analysis. Proteins extraction and western blot analysis were performed as previously described^66^.

### Ethylene emanation

Hundred seedlings of Arabidopsis Col-0 were cultivated *in vitro* during 14 days in square plates (12 x 12 cm) as described in^67^. The measurements were then performed over a continuative period of 24 hours for each plate.

## Statistics and Reproducibility

Data shown in graphs are presented either as the mean with S.D. or as median values from three biological replicates as indicated in the figure captions. Statistical significance was tested by one-way ANOVA test to compare treatments with mock (asterisks denotate significant difference, p < 0.05) whereas one-way ANOVA test followed by Tukey’s post-hoc test was performed to compare different treatments to each other and mock (different letters indicate significant differences, p< 0.05). Students’ t-test (p < 0.05) was used to identify differentially expressed genes from transcriptome data.

## Data Availability statement

Microarray transcriptomic data generated in the current study have been submitted to the ArrayExpress repository, accession E-MTAB-10192. All other source data are included in the article as supplementary raw data.

## Abbreviations

AA: auxiliary activity
AA9_COS: AA9-derived cello-oligosaccharides or cellulose-oligosaccharides
COS: cello-oligosaccharides
CWDE: cell wall degrading enzyme
CWI: cell wall integrity
DAMP: damage-associated molecular patterns
DP: degree of polymerization
ET: ethylene
ETI: effector triggered immunity
GH: glycoside hydrolase
JA: jasmonic acid
LPMO: lytic polysaccharide monooxygenase
LRR-RLK: leucine-rich repeat receptor-like kinases
MAPK: mitogen-activated protein kinase
OG: oligogalacturonide
PAMP: pathogen-associated molecular-pattern
PASC: phosphoric acid swollen cellulose
PTI: pattern triggered immunity
DTI: damage triggered immunity
SA: salicylic acid

## Acknowledgments

This work was supported by FNRS-MIS LUX-project F.4502.19 starting grant to DC (MZ and AM) and PINT-BILAT-M R.M012.18; also by INNOVIRIS – 2019-Bridge-4: Re4Bru (MASK, SM) for productions of LPMOs, FER-2017 for the HPAEC-PAD analytical platform. We thank Prof. Cédric Delporte of the Analytical Platform of the Faculty of Pharmacy (APFP), ULB for the mass spectrometry of camalexin. APFP is supported by Belgian National Fund for Scientific Research (FRS-FNRS) and ULB funds (FRS N° 3.4553.08 and T.0136.13; ULB FER-2007; FER-2013; FER-2019 and Platform funds). We also thanks Dr. Laurent Grumiau and Dr. Florence Rodriguez (ULB, molecular biology platform) for technical assistance during immunoblotting analysis. MC is a Research Associate of the INRAE (France). The IJPB benefits from the support of Saclay Plant Sciences-SPS (ANR-17-EUR-0007). This work has benefited from the support of IJPB’s Plant Observatory technological platforms, Dr. G. Mouille for SA and JA mass spectrometry analysis. We are grateful to Dr. Marie-Christine Soulié for providing *B. cinerea* B05.10 strain and Dr. Herman Höfte for providing us *BAK1* and *THE1* loss-of-function lines, all from IJPB-INRAE-Versailles, FR. Finally, the authors are deeply grateful to the reviewers for providing exceptional meaningful revision suggesting experiments that markedly improved the final output of the work.

## Author Contributions

MZ and DC conceptualized, planned and executed experiments, plant growth and treatments. MZ performed transcriptome, plant physiology, phytopathology, immunoblotting tests and analytics. MZ, AM and IM performed the qRT-PCR analyses. MASK, SM, MOG, AM, and DC cloned the enzymes, performed HPAEC and LPMO biochemistry. MZ and MC performed bioinformatics, GO, and network analysis. MZ, SJ and MF performed metabolite extractions and callose staining. MZ and IM performed western blot analysis. MZ and DC wrote the paper with contributions from all the coauthors. DC conceptualized and supervised the work; obtained grants, set analytical equipment, established plant growth conditions and enzymatic cloning platform. All authors read and approved the manuscript for publication.

